# Limited plasticity in gene expression for providing or receiving parental care under different temperatures

**DOI:** 10.1101/2021.12.14.472162

**Authors:** Jeanette B. Moss, Christopher B. Cunningham, Elizabeth C. McKinney, Allen J. Moore

## Abstract

Parenting is thought to evolve to buffer offspring from variable, unpredictable, and challenging environmental conditions. In the subsocial carrion beetle, *Nicrophorus orbicollis*, stressful temperatures during parenting do not affect parental behavior despite imposing steep fitness costs to parents. Here, we ask if plasticity of gene expression underpins this behavioral stability or facilitates independent compensation by larvae. To test this we characterized gene expression of parents and offspring before and during active parenting under benign (20°C) and stressful (24°C) temperatures to identify genes of parents and offspring associated with thermal response, parenting/being parented, and gene expression plasticity associated with behavioral stability of parental care. The main effects of thermal and social condition each shaped patterns of gene expression in females, males, and larvae. In addition, we implicated 79 genes in females as ‘buffering’ parental behavior across environments. The majority of these underwent significant changes in expression in actively parenting mothers at the benign temperature, but not at the stressful temperature. Our results suggest that neither genetic programs for parenting nor their effects on offspring gene expression are fundamentally different under stressful conditions, and that behavioral stability is associated primarily with the maintenance of existing genetic programs rather than replacement or supplementation. Thus, while selection for compensatory gene expression could expand the range of thermal conditions parents will tolerate, without expanding the toolkit of genes involved selection is unlikely to lead to adaptive changes of function.

## INTRODUCTION

Parental care evolves to buffer offspring against environmental stress and once evolved, can be instrumental for ensuring success across a range of ecological conditions even beyond those that promoted its evolution (Clutton-Brock, 1991; Royle, Smiseth, & Kölliker, 2012; Wilson, 1975). In the face of rapidly changing environments, phenotypic plasticity in family interactions could contribute to species persistence by allowing parents to invest more when conditions are poor and for offspring to compensate when conditions exceed those that parents can or are willing to buffer. In some cases, this phenotypic plasticity may be immediately detectable as a change in behavior. For example, many birds compensate for hot breeding season conditions by extending their time and effort at the nest, sometimes risking dehydration to maintain stable incubation conditions for embryos (McClintock, Hepp, & Kennamer, 2014; Clauser & McRae, 2016; Sharpe, Bayter, & Gardner, 2021), while offspring have been shown to upregulate metabolic and immune pathways in response to the removal of a caring parent (Körner et al., 2020; Mashoodh, Westoby, & Kilner, 2021; Ziadie, Ebot-Ojong, McKinney, & Moore, 2019) or exposure to pathogens (Körner et al., 2020). However, when parents and offspring are pushed to their physiological limits, maintaining a phenotype stability (i.e., continuing to provide care at the same level) is likely just as important for ensuring offspring success as is altering behavior.

Transcriptomic approaches are useful for examining gene expression plasticity beneath the surface of phenotypes because gene expression is inherently plastic, and gene expression profiles can undergo profound changes while maintaining stable higher-level phenotypes (Eberwine & Kim, 2015; Nijhout, Best, & Reed, 2019; Rivera et al., 2021); that is, many ‘genetic programs’ can give rise to identical phenotypes. While most research in this area has focused on the role of plastic gene expression in maintaining physiological performance and stabilizing morphological traits under stress (Oleksiak, Roach, & Crawford, 2005; Cheviron, Whitehead, & Brumfield, 2008; Cheviron & Brumfield, 2011; Peck et al. 2015), compensatory molecular mechanisms are also expected to underpin stability of complex behaviors across fluctuating environments (Fischer, Hauber, & Bell, 2021). This perspective is valuable to the study of parental care because parenting evolves to minimize environmental fluctuations and involves transformational changes of gene expression (Parker et al. 2015; Ray et al. 2016; Bukhari et a. 2019; Fischer et al. 2019; Lopes & de Brujn), and the ultimate configuration of expressed genes associated with care may differ depending on environment. Because parental care is inherently a social exchange and phenotypic plasticity can stem from donors and/or recipients of care, a comprehensive test of these predictions should profile the gene expression of parents and offspring simultaneously (Butler & Maruska 2021).

Here, we performed RNA-seq on a subsocial burying beetle, *Nicrophorus orbicollis*, to explore if plasticity of gene expression helps buffer families against environmental stressors. There is a strong basis for studying the genetics of parenting and development of burying beetles (Benowitz, McKinney, Cunningham, & Moore, 2017; Cunningham et al., 2019; Jacobs et al., 2016; Palmer et al., 2016; Parker et al., 2015; Won et al., 2018), yet the genes responsible for mounting responses to major environmental stressors while beetles are parenting/being parented remain uncharacterized. Such mechanisms are particularly relevant in the context of rapid environmental change, as they could provide targets of selection in populations unable to behaviorally avoid stressful conditions. Even less understood is the extent to which application of a secondary stressor may modify core parental responses; i.e., how plasticity in behavioral gene expression buffers higher level phenotypes.

Higher temperatures during parenting for *N. orbicollis* impose steep reproductive and survival costs; however, females and males do not change their parental care behavior, type, or intensity (Moss and Moore, 2021). Whether behavioral stability of parents results from gene expression changes that alter or supplement the genetic program for parental care, or whether gene expression changes in offspring may independently compensate for thermal stress, is not known. Here, we address these possibilities while controlling for any confounding effects of differences in care. We examined the effects of two thermal environments to characterize ‘thermal response’ genes (20°C or 24°C, the same as that investigated in Moss and Moore, 2021) and parent-offspring interactions to characterize ‘parenting/being parented’ genes (before or during active parenting), comparing responses of mothers, fathers, and offspring to understand the similarity of each’s response. Our prediction was that thermal stressors and parental care will each elicit independent gene expression responses and that these will differ between family members (represented by the main effects of the statistical model). Further, we predicted that some genes would show distinct patterns of expression change in response to the combination of stressors to stabilize the behaviors across environments (“buffering genes” represented by the interaction term of the statistical model); specifically, higher temperatures should significantly modify expression of core genes for parenting/being parented or elicit distinct changes in gene expression during family interactions if genetic plasticity facilitates behavioral stability across environments.

## MATERIALS AND METHODS

### Study System

*Nicrophorus orbicollis* is a biparental carrion beetle that breeds on small vertebrate carcasses in woodlands ranging from southern Canada to northern Texas. At the southeastern edge of this distribution, daily mean temperatures as low as 18°C and as high as 25°C may arise over the course of the long summer breeding season (Moss & Moore, 2021). Temperatures at the upper end of this range (23–25°C) are highly challenging for parents and developing offspring, with families forced to breed under these conditions suffering shortened adult and larval lifespans and reduced clutch size larval mass (Moss & Moore, 2021). As in most members of the genus, *N. orbicollis* expresses a large repertoire of pre- and post-hatching care behaviors (Eggert & Müller, 1997; Scott, 1998). Carcass preparation begins with the removal of hair and liberal application of anal excretions, which suppress microbial growth on the prepared ‘brood ball.’ Females oviposit throughout the soil surrounding the brood ball, and within 2–3 days these eggs hatch and larvae migrate through the soil to the brood ball attended by their parents. At this stage, parents commence directly provisioning begging young *via* regurgitation. While larvae of *N. orbicollis* have relatively high starvation tolerance (mean time to starvation: ∼17 hours), upon hatching their capacity to self-feed from a prepared carcass is extremely limited, and they rely on at least three hours of post-hatching parental care for survival (Capodeanu-Nägler et al. 2018). This post-hatching stage is likely to be the most energetically demanding stage of care for parents as it is when attendance to offspring is highest and the stage at which we see the most sex differences of parental behavior (Moss & Moore, 2021).

### Study Design

We used *N. orbicollis* beetles that had been collected locally and reared in the laboratory for one generation at the University of Georgia, as described in Moss and Moore (2021). Different beetles were used to generate behavioral (Moss and Moore 2021) and transcriptomic datasets (this study). Briefly, adults were housed individually in plastic containers (9 cm diameter, 4 cm deep; Eco Products, Boulder, CO, USA) filled with potting soil and fed organic ground beef twice weekly *ad libitum*. Stock breeding took place at room temperature (20 ± 0.5°C), and resulting offspring were divided evenly between two incubators starting three days into pupation for a thermal acclimation period prior to breeding. These incubators were programmed to long-day light conditions (reverse light:dark 14:10) on ramping temperature cycles to simulate a diurnal range: the first fluctuating between 21 and 20°C and the second between 25 and 24°C. Because breeding takes place underground in relatively thermostable conditions, however, only the lower end of the diurnal range of each thermal treatment (20°C and 24°C) was used for breeding. Virgin, non-sibling beetles (aged ≥ 14 days post-eclosion) were paired in plastic boxes (17.2 × 12.7 × 6.4 cm; Pioneer Plastics, Dixon, KY, USA) filled with approximately 2 cm of moistened soil and a 40–45 g thawed mouse carcass (RodentPro, Evansville, IN, USA) and moved to either a constant dark temperature-controlled room (20°C) or incubator (24°C) corresponding to their acclimation environment. Boxes were monitored twice daily for the presence of eggs, and after 2–3 days eggs were collected into petri dishes containing damp filter paper and stored at their respective treatment temperatures. Eggs were checked every four hours between 0800 and 1700 until hatching, which occurs within two days after laying.

In addition to manipulating the thermal environments in which care took place, we also generated within-temperature treatment controls to disentangle the effects of “active” parenting (the stage at which parents and larvae interact socially, which we refer to as parenting or direct parental care) from a general parenting state which includes behavior such as carcass preparation that does not require social interactions and begins before larvae are present. Thus, we sampled at two time points: 1) post-larval hatching but before larvae and parents interacted (≤ 16 hours old larvae); and 2) after the first 24 hours of direct parental care. These larvae were collected directly from the petri dish immediately when found to have hatched, with parents collected simultaneously from the carcass they were preparing. Larvae of families for parenting samples, by contrast, were transferred to brood balls attended by parents when found to have hatched. These families were left undisturbed for 24 hours so that direct parenting could take place before collecting both parents and offspring (24–40 hours old larvae) for analysis. Parents in both treatments had initiated indirect parental care involving preparation of a brood ball for larval feeding. Timing of sampling parents before interactions coincided with the circadian switch to active parenting (Oldekop, Smiseth, Piggins, & Moore, 2007).

### Sample Collection, Preparation, and RNA Sequencing

Our design yielded ten families at 20°C (five before parenting, five during parenting) and nine families at 24°C (four before parenting, five during parenting). We collected the heads of adults, consisting of brain and associated “fat body” tissue, as in previous studies (Parker et al 2015). This allowed us to capture gene expression in the two tissue types that influence behavior in insects. Because of the size of larvae at hatching and because we had no *a-priori* predictions for the specific tissues affected by interactions with adults, we collected whole bodies. Larval samples therefore consisted of seven to ten larvae for families before parenting and two to four larvae for families during parenting. Individual adult samples, both before and during parenting, were age-matched. Samples were flash frozen in liquid nitrogen and stored at -80°C until RNA extraction. Extractions were performed following the Qiagen RNAeasy Lipid kit protocols (Qiagen, Venlo, the Netherlands). Sequencing libraries were prepared from 1.3 μg total RNA using the Illumina TruSeq mRNA Stranded Library Kit (Illumina, Inc., San Diego, CA, USA) according to the manufacturer’s protocol with standard Illumina adapters and primers. Sequencing was carried out on a NovaSeq 6000 platform with a 150 bp paired-end protocol, targeting 40M read pairs per sample.

### Read Mapping, Transcriptome Assembly, and Annotation

Reads were trimmed using Trimmomatic (Version 0.39; Bolger et al. 2014) to remove adapter sequences with default parameters, except low-quality bases (Phred <15 at the leading and trailing ends and from 4-base sliding windows), and short reads (<36 bp). Reads were corrected using RCorrector with default parameters (Version 1.0.3.1; Song and Florea 2015). Quality was assessed using FastQC (Version 0.11.9; http://www.bioinformatics.babraham.ac.uk/projects/fastqc/; Supplementary Table 1). Paired-end reads were mapped against a draft reassembly of the WGS data of Benowitz et al. (2017) of *N. orbicollis* genome (NCBI BioProject PRJNA371654; Supplementary Methods 1). Read mapping was performed using HISAT2 with default parameters (Version 2.1.0; Kim et al. 2015). A reference-guided transcriptome was assembled using StringTie (Version 2.1.1; Pertea et al. 2015) and redundant transcripts were removed using CD-HIT (Version 4.8.1; Li and Godzik 2006) with a sequence similarity cut-off value of 95%. Finally, we assessed the completeness of the assembly using BUSCO (Insecta dataset; Version 5.0.0; Simão et al. 2015).

Annotation of contigs was performed using a combination of approaches. Briefly, the longest isoform of each gene was screened against the annotated and published genome of the closely related burying beetle *N. vespilloides* (NCBI Assembly Accession No. GCF_001412225.1; Cunningham et al., 2015) using Magic-BLAST (Version 1.5.0; Boratyn et al. 2019). Transcripts were also fed through the ‘dammit’ pipeline (http://dib-lab.github.io/dammit/), which identifies candidate coding regions in transcripts with Transdecoder (Version 2.0.1; https://hpc.ilri.cgiar.org/transdecoder-software) and annotates them, drawing from multiple databases. We used the *N. vespilloides* proteome (Cunningham et al., 2015) as a user database to search against and also retained any hits to known protein domains in Pfam-A (Version 32.0; El-Gebali et al. 2019) and orthologs in OrthoDB (Version 10; Kriventseva et al. 2019). Orthologous genes were retrieved from the OrthoDB database by searching cluster IDs at the level Polyphaga. Gene ontology (GO) terms for annotated genes were assigned by using Pfam accession numbers and GO terms of OrthoDB assignment. GO terms were also supplemented using eggNOG-mapper (Version 5.0; Huerta-Cepas et al. 2019) and OrthoDB searches at the Order (Insecta) level. Unique GO terms from any source were retained for a gene.

### Transcript Quantification and Filtering

We estimated transcript abundances for each sample separately using StringTie. A transcript count matrix was extracted with the prepDE.py script provided with StringTie and imported into R (Version 4.0.3; R Core Development Team 2019) along with sample library sizes (calculated using Picardtools, Version 2.21.6; http://picard.sourceforge.net) and transcript lengths. We computed the fragments per kilobase per million (FPKM) matrix with the package *countToFPKM* (Version 1.0; Alhendi 2019). For each family member and social state (before or during parenting), we retained transcripts that were expressed >2 FPKM in more than half of samples and then combined all unique transcripts across groups in downstream analyses (Bloch et al., 2018). Finally, we used *IsoformSwitchAnalyzeR* (Version 1.12.0; Vitting-Seerup and Sandelin 2019) to convert transcript-level counts into gene-level counts.

### Differential Expression Analysis

We performed differential expression analysis using *DESeq2* (Bioconductor Version 1.30.0; Love et al. 2014). To cluster samples, we first applied variance stabilizing transformation using DESeq2 and performed a principal components analysis (PCA) using R’s built-in ‘prcomp’ function. Visualization of the clustering was done using *DESeq2*’s ‘pcaplot’ function and extracted principal components were regressed on each grouping factor (family member, temperature, parenting, and the statistical interactions among these factors) *via* analysis of variance (ANOVA using R). Significant differences among family members in the full model ANOVA led us to analyze males, females, and larvae separately. For each family member, differential expression was estimated using parametric dispersion. We used the likelihood ratio test (LRT) implemented in *DESeq2* to compare the goodness-of-fit of several models for each family member to estimate differential gene expression. Two tests were used to capture gene expression changes in response to main effects. The first compared a full model fitted with both temperature and parenting (temperature + parenting) to a reduced model fitted only with parenting to identify differentially expressed genes mutually affected by temperature across parenting states (henceforth, ‘thermal response’ genes). The second compared the same full model to a reduced model fitted only with temperature to produce a list of differentially expressed genes mutually affected by parenting across thermal environments (henceforth, ‘parenting’ genes). The *p-*values that resulted from these tests were adjusted for multiple testing using the Benjamini-Hochberg correction (Benjamini & Hochberg, 1995) and genes with adjusted *p* < 0.05 were considered statistically significantly differentially expressed genes.

To assess similarity of gene expression responses to the main effects of temperature and parenting across family members, we extracted lists of overlapping genes from the intersect of male, female, and larval parenting/being parented and thermal response gene sets. To determine whether overlap was significantly greater than expected by chance given variable input sizes, we further carried out randomization tests. Briefly, 10,000 gene sets were randomly generated for males, females, and larvae by sampling from the full list of expressed genes in that family member a number equivalent to the size of the gene set being simulated. These random gene sets were used to calculate a null distribution of overlap values, and significance was calculated as the proportion exceeding the observed overlap for each pairwise comparison. While overlap of the most highly significant genes between two sets lends itself to intuitive interpretation, use of fixed thresholds can be overly stringent and lead to false negatives. Therefore, we implemented a complementary threshold-free approach: rank-rank hypergeometric overlap (RRHO) analysis. This analysis ranks DE genes according to their significance level along with the magnitude and direction of change to make qualitative assessments of concordance between complete gene sets independent of *p*-value cutoff. Results were visualized using the stratified method of the R package *RRHO2* (Version 1.0; Cahill et al. 2018), with genes ranked according to the recommended metric of -log10(*p*-value)*sign(log_2_ fold change).

Finally, we assessed whether the genetic program for parenting/being parented was altered or supplemented under the addition of thermal stress in two ways. First, we performed a differential expression analysis using the LRT approach to identify genes showing distinct responses to parenting/being parented (substantial differences in magnitude or different signs) in the presence of thermal stress (henceforth, ‘buffering’ genes). To obtain this list, we compared a full model fitted with a statistical interaction term (temperature x parenting) to a reduced model fitted only with the main effects (temperature + parenting). To further characterize patterns of differential gene expression across social and temperature groups, we performed an ANOVA followed by post-hoc contrasts (Tukey HSD) at the level of each gene in the buffering set, comparing the magnitude of expression and sign of the response at 24°C vs 20°C. Second, we compared differential expression of parenting/being parented genes between the two thermal environments. Even if changes of expression of these genes are similar enough across thermal environments to be captured as main effects, the magnitude of the response may be enhanced or dampened due to thermal stress. To test this possibility, we estimated log2fold changes of expression of parenting/being parented genes separately for each temperature treatment using the ‘ashr’ shrinkage estimator with false discovery rate correction (Stephens 2017) and fit a regression to these data. A slope different than one allows rejection of the null hypothesis that responses are generally similar in magnitude across thermal treatments and provides a more holistic comparison of the treatments.

### Functional Enrichment Analysis and Candidate Genes

We queried the differentially expressed gene sets and the overlapping gene sets for enriched Gene Ontology terms using topGO (Version 2.42.0; Alexa and Rahnenfuhrer 2020) searching for enrichment in all three gene ontology categories: biological process, molecular function, and cellular component. The nodeSize parameter was set to 10 to remove GO terms with fewer than ten annotated genes and only terms with >1 significant gene were retained. Significance was estimated using a Fisher’s exact test following the program’s suggested protocol. To examine the directionality of change of enriched pathways, Z-scores were calculated following the formula of Walter, Sánchez-Cabo, & Ricote (2015): number of downregulated genes annotated subtracted from the number of upregulated genes annotated for the term divided by the square-root of the total number of genes annotated for the term.

## RESULTS

Our final transcriptome assembly contained 91% of single-copy conserved nucleotide orthologs (Complete: 90.5% [Single Copy: 76.4%, Duplicated: 14.1%], Fragmented: 4.2%, Missing: 5.3%) and a total of 12,406 genes.

### Thermal and Social Conditions Shape Gene Expression of Family Members

We visualized differences in gene expression among male parents, female parents, and offspring using principal components analysis. Parents and offspring separated strongly along two primary axes (Fig 1A), which together explained 77.8% of total variance in gene expression across all samples (PC1: 73.08%; PC2: 4.76%; Table 1). Family member (parent vs larvae) clearly separated along PC1, while separation along PC2 was driven primarily by parenting (Table 1). The interactive effect of family member and parenting and family member and temperature were significantly associated with PC2 (Table 1; Fig 1B). Visualizing family members separately with their own dispersions further clarified how global gene expression differed among groups. Female gene expression displayed considerable overlap between thermal environments but was most divided with respect to parenting, particularly in the 20°C environment (Fig 1C). Male gene expression differentiated most strongly with temperature (Fig 1D). In larvae, behavioral groups formed distinct clusters, with parenting accounting for far greater among-group differences than temperature (Fig 1E).

**Figure 1:**
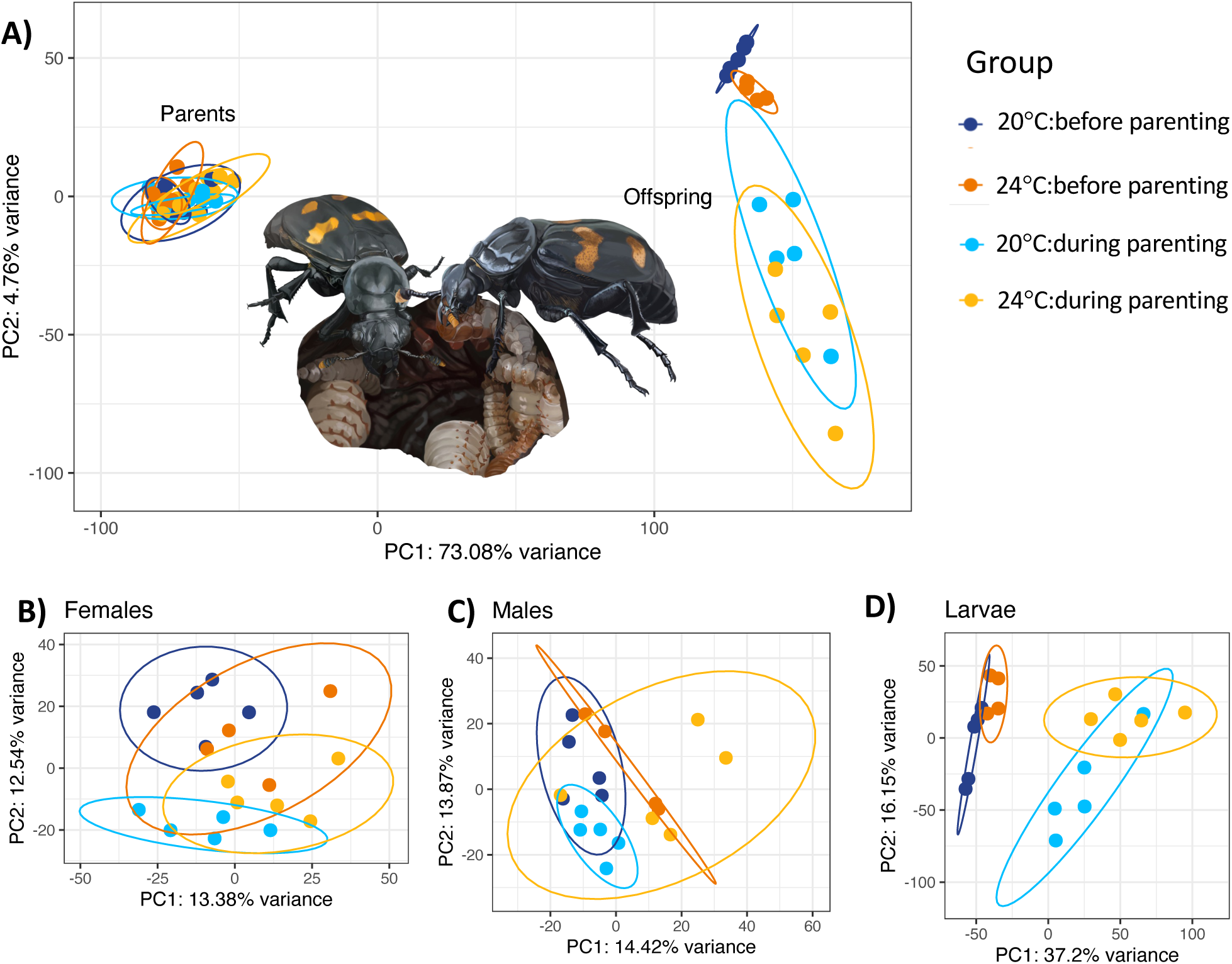
Principal components analysis (PCA) of normalized transcriptomic profiles across the three family members (females, males, and larvae) and four treatment groups (20°C:before parenting, 20°C:during parenting, 24°C:before parenting, and 24°C:during parenting). (A) Sample clustering along the first two principal components of the full model PCA (∼Family Member*Temperature*Parenting). Eclipses are drawn for 95% confidence intervals. (B–D) Clustering of female, male, and larval samples, respectively, when analysis is repeated using only those samples. Illustration of *N. orbicollis* family in Fig 1A by K.E. Kollars.

**Table 1:** ANOVA of principal components of overall gene expression.

| Variable | PC1 (73.08% variance) |  | PC2 (4.76% variance) |  |
| --- | --- | --- | --- | --- |
|  | F<sub>2,55</sub> | *P* | F<sub>2,55</sub> | *P* |
| Family Member | **5346.224** | <0.0001 | 0.624 | 0.540 |
| Temp | **4.047** | 0.050 | **7.073** | 0.011 |
| Parenting | **27.238** | <0.0001 | **88.311** | <0.0001 |
| Family Member x Temperature | 0.499 | 0.611 | **11.406** | <0.0001 |
| Family Member x Parenting | **4.852** | 0.012 | **92.516** | <0.0001 |
| Temperature x Parenting | 2.139 | 0.151 | 0.925 | 0.341 |
| Family Member x Temperature x Parenting | 1.653 | 0.203 | 1.287 | 0.286 |

### Plasticity of Gene Expression Underpins Thermal Response, Parenting/Being Parented, and Buffering

We used differential expression analysis to characterize plastic changes in gene expression associated with the main effects of temperature and parenting, as well as their interaction. We identified genes showing both shared (overlapping) and distinct responses to main effects across family members, consistent with variation in physiology and behavior between sexes and life stages. There was also gene expression variation associated with buffering of behavioral responses, where family members expressing the same behaviors in different environments showed different patterns of gene expression. These included subtle expression variation of parenting/being parented genes across temperature treatments, as well as some changes that occurred specifically in response to the interaction of temperature and parenting.

#### i. Thermal Response Genes

Temperature induced a stronger gene expression response in males (n = 418 differentially expressed genes; Supplementary Table 2) and larvae (n = 487 differentially expressed genes; Supplementary Table 3) than in females (n = 284 differentially expressed genes; Supplementary Table 4). However, male, female, and larval responses to temperature showed statistically significant overlap in terms of gene identity (male-female: *P* < 0.001; female-larvae: *P* < 0.001; male-larvae: *P* = 0.006; Fig 2A). Global concordance was pronounced between males and females (Fig 2C) – both in genes that were up-regulated and down-regulated in response to high temperature – whereas this signal was weak when comparing females to larvae (Fig 2D) and males to larvae (Fig 2E).

**Figure 2:**
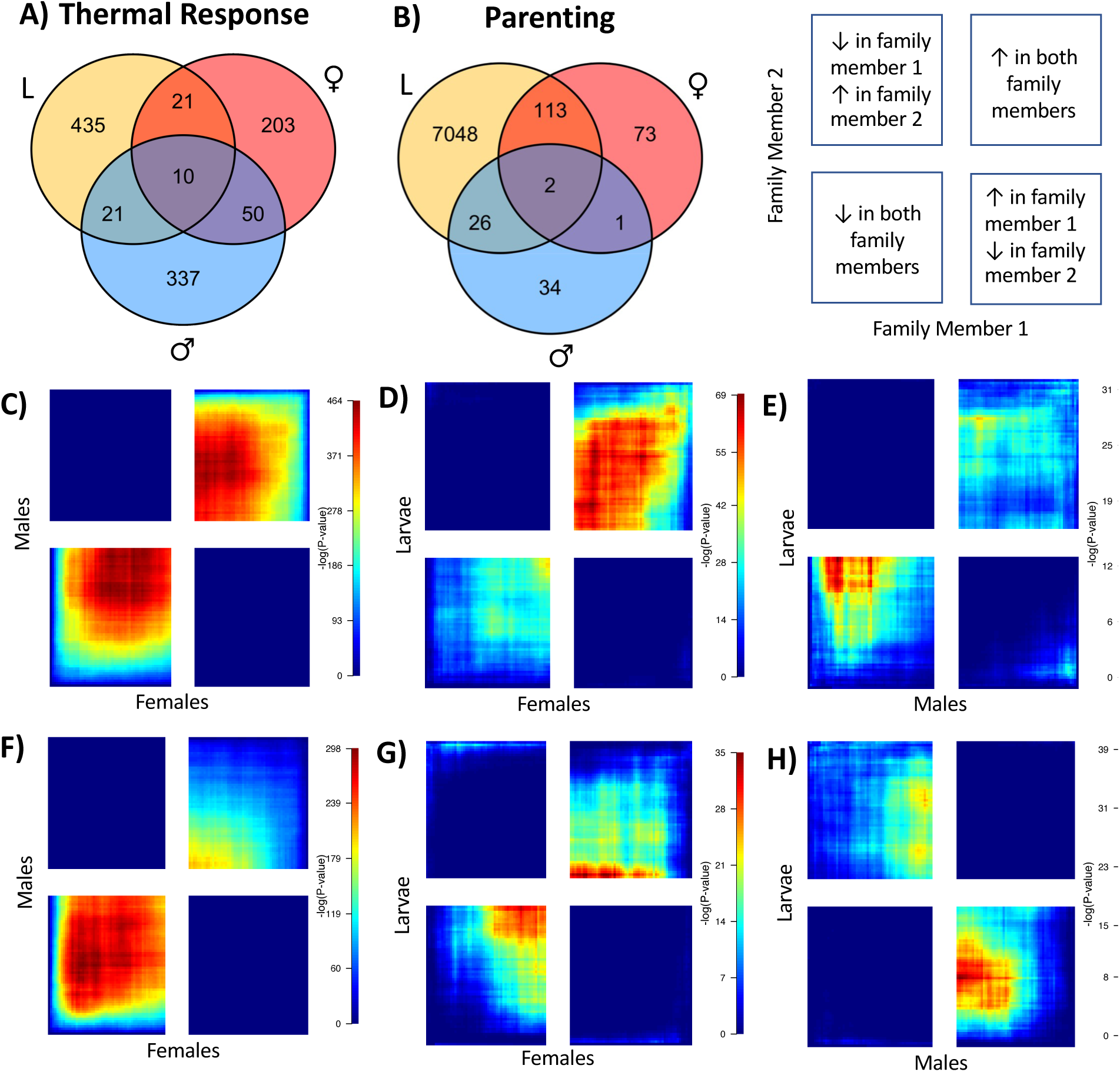
Concordance in differential gene expression between family members as estimated by direct overlap of differentially expressed genes (A–B) and rank-rank hypergeometric overlap (RRHO; C–H). Numbers of shared differentially expressed genes associated with (A) thermal response and (B) parenting are shown in Venn diagrams for females (red), males (blue), and larvae (yellow). The bottom panel shows the results of the RRHO analysis of all 12,406 genes for each of the respective contrasts (20°C vs 24°C: C–E; and with vs without parenting: F–H). The x (family member 1) and y (family member 2) axes of each plot correspond to the significance of differential expression (ranked -log10(*p*-values)) multiplied by the signed log2fold difference in gene expression for each family member-specific analysis. White boundaries demarcate the switch from down-to up-regulation between behavioral groups for each family member. Pixel color corresponds to the value of the -log10(*p*-value) from the differential expression analysis, such that hot spots in the plot designate the strength and directionality of concordance in gene expression between family members.

#### ii. Parenting Genes

With the parenting main effect, the highest number of differentially expressed genes was associated with being parented, with 7,189 genes differentiating larvae before and during interactions with parents (Supplementary Table 5). Adults had fewer differentially expressed genes than larvae, and within adults there were sex differences. Fewer differentially expressed genes were detected in males after the arrival of larvae (n = 63; Supplementary Table 6) than in females (n =189; Supplementary Table 7). Despite this, females and males showed statistically significant overlap of gene identities while parenting (Fig 2B; *P* = 0.044). When considering all expressed genes, concordance was striking but directional, with overlap between the sexes concentrated in genes downregulated in both during larval interaction (Fig 2F). Females interacting with larvae and larvae interacting with parents shared more differentially expressed genes in common (n = 115) than parenting males and parented larvae (n = 28), although this did not reach statistical significance (*P* = 0.059) and neither comparison showed global signatures of concordance of gene expression (Fig 2G–H).

#### iii. Buffering Genes

To evaluate the role of gene expression plasticity in facilitating parental buffering and offspring compensation under stressful environments, we performed a transcriptome-wide survey for genes showing distinct responses to parenting/being parented at 24°C versus 20°C, or a significant interaction between temperature and parenting. More genes met this criterion and were differentially expressed in females (n = 79; Supplementary Table 8) than in males (n = 4; Supplementary Table 9) or larvae (n = 26; Supplementary Table 10). To test whether genes showing plasticity indeed showed stronger and broader expression changes at the more stressful temperature, the 79 buffering genes in females were subjected to closer examination (Supplementary Table 11). At the benign temperature (20°C), most genes (n = 57) showed increases of expression under active parenting, whereas at the higher temperature (24°C) only five genes showed any change in expression. Hence, most buffering genes fell into two categories: gene expression levels before parenting were equivalent between thermal treatments (n = 35) or gene expression levels started out significantly higher at 24°C than at 20°C (n = 32; e.g., Apolipophorin-III; Fig 3C). Because plasticity was temperature-dependent, gene expression levels during parenting were either significantly lower at 24°C (n = 29) or did not differ between thermal environments (n = 45), and only five genes showed significantly higher expression levels during parenting in the warmer treatment.

**Figure 3:**
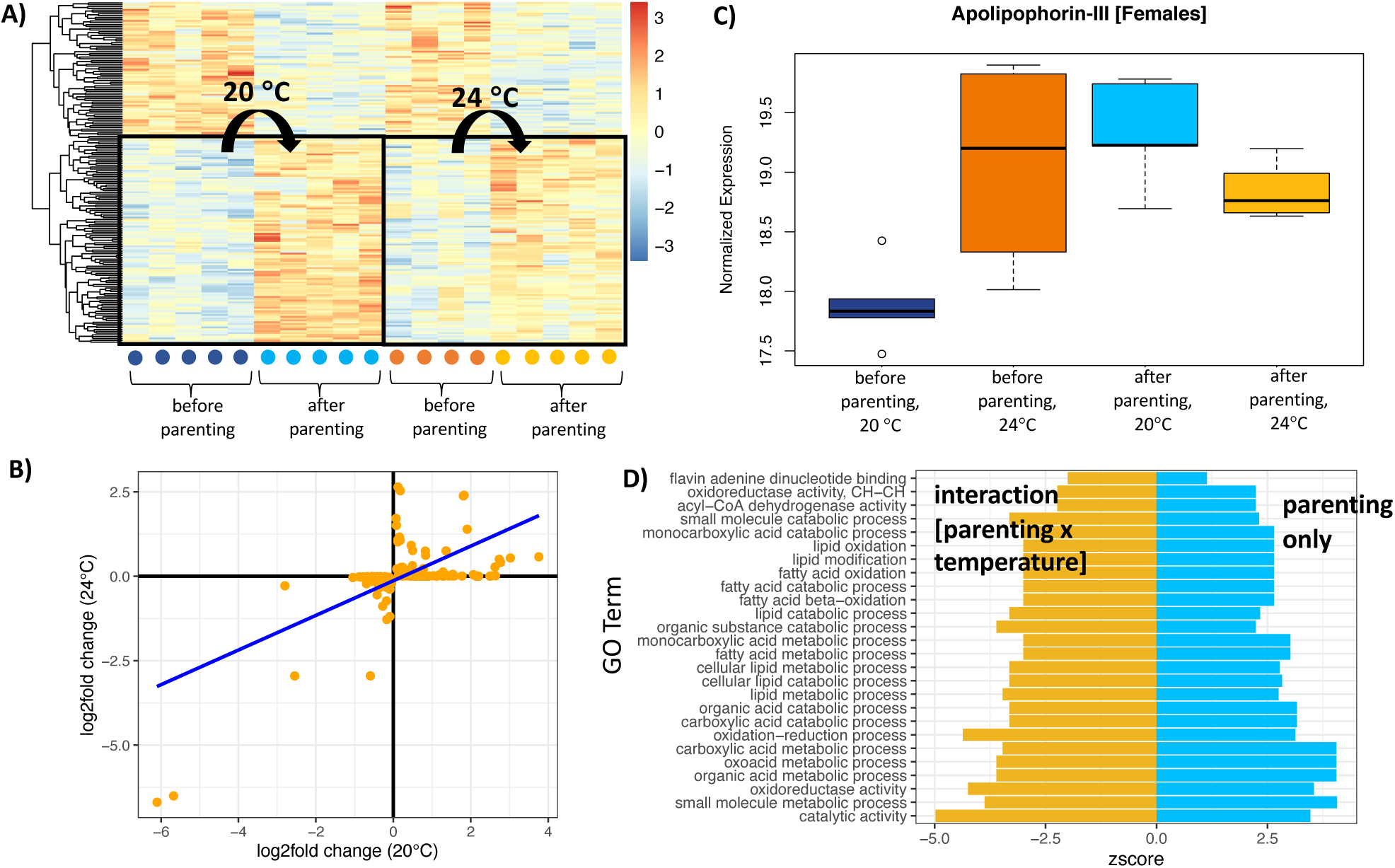
(A) Normalized expression profiles of the 189 genes that were differentially expressed in females before and during parenting depicted as heatmaps, with each column representing an individual sample (grouped by social and temperature treatment) and each row representing a gene (clustered in a dendrogram by expression similarity). Cell colors represent normalized count values, such that the visual distinctiveness of clusters of samples corresponds to the magnitude of difference in their gene expression between groups; (B) Regression of log2fold changes in expression of 189 genes that were differentially expressed in females before and during parenting between 24°C and 20°C environments; (C) Distribution of normalized expression values of Apolipophorin-III across four behavioral groups of females: before parenting at 20°C, before parenting at 24°C, during parenting at 20°C, and during parenting at 24°C; (D) Directionality of expression (calculated as a z-score) of 26 GO terms identified as significantly enriched in females in response to parenting only (blue bars) as well as in response to the interaction of parenting and temperature (gold bars). Z-scores correspond to the standardized number of genes annotated for a term that are upregulated relative to downregulated, such that values less than zero correspond to overall downregulation.

We next compared gene expression responses to parenting/being parented across temperature treatments. Among the core genes for parenting/being parented (genes that were differentially expressed between parenting states across temperatures), differential expression was weaker at 24°C relative to 20°C in females (slope = 0.514; Fig3A, B) and males (slope = 0.357), whereas the slope of this regression in larvae was close to one (slope = 0.976), suggesting that responses of larvae were generally similar in magnitude across thermal environments.

### Functional Enrichment Analysis

Functional enrichment of GO terms in the within-family member temperature contrasts revealed both shared and distinct enriched functional characteristics across females, males, and larvae. The most significantly enriched GO terms in parents were related to chitin and carbohydrate metabolism (Supplementary Table 12–13), whereas larvae showed the most enrichment for cellular housekeeping functions (e.g. ribosome biogenesis and rRNA processing; Supplementary Table 14). Compared to the 20°C group, females, males, and larvae in the 24°C group predominantly showed downregulation of functional pathways.

Actively parenting females showed significant positive enrichment in a range of metabolic, catabolic, and biosynthetic functions compared to their respective non-interacting controls (Supplementary Table 15). Forty-five of the 64 GO terms that were enriched in the female parenting gene set were also significantly enriched in the overlap between female parenting genes and larval parented genes, particularly terms related to the metabolism of organic, carboxylic, and fatty acids (Supplementary Table 16). Conversely, analysis of GO terms in males revealed significant downregulation of non-overlapping GO terms (e.g. larval cuticle patterning; Supplementary Table 17). Larval functional pathways were predominantly upregulated in response to interactions with parents and spanned many categories, from cellular organization and differentiation to whole organism development (Supplementary Table 18).

Finally, we examined Gene Ontology term enrichment under the interaction of parenting and thermal environment. Buffering genes of males (n = 4) showed enrichment in three pathways related to lipid transport and localization (Supplementary Table 20) while those in larvae (n = 26) showed enrichment in five pathways (Supplementary Table 21), the most significant of which (synaptic target inhibition and negative chemotaxis) were upregulated in parented larvae responding to thermal stress. In females interacting with larvae, we found 45 significantly enriched GO term annotations for genes that were differentially expressed between 20°C and 24°C, including 26 GO terms that were also enriched in the female parenting gene set but in the opposite direction (Supplementary Tables 12, 19; Fig 3D). Top GO terms related to the oxidation and catabolism of fatty and organic acids and comprised genes that were significantly downregulated, on average (Supplementary Table 19).

## DISCUSSION

Parenting is expected to evolve to buffer environmental stress, but parenting involves an interaction between the parent and the offspring. For parenting to be effective, the family interaction needs to be predictable and the mechanisms responsible for stabilizing these interactions across variable environments are not well understood. Plasticity of gene expression could allow families to optimize dynamics for their current environment. If true, then parenting-related responses to an ecologically relevant stressor should have a gene expression signature. We test this prediction in a subsocial insect with biparental care. We found that thermal and social conditions both shape patterns of gene expression in *N. orbicollis* females, males, and larvae, and that many responses are distinct to specific family members. Further, we found evidence for some plasticity of gene expression in response to the combination of social and thermal stressors. Specifically, parenting males and females, but not parented larvae, showed weaker gene expression responses to interactions with larvae under stressful temperatures. In females 79 genes responded to parenting in an entirely temperature-dependent manner, showing significant changes in expression at the benign temperature but not at the stressful temperature. Taken together with our previous findings of limited behavioral plasticity of *N. orbicollis* parents exposed to stressful temperatures (Moss & Moore, 2021), this implies that behavioral stability under stress is associated primarily with maintenance of existing genetic programs, rather than switching to an alternative or supplemented genetic program for the same complex behavior.

Parental investment and offspring development are sensitive to environmental quality as poor conditions can impose competing physiological demands between parenting and thermal response. To capture and compare the gene expression and hence the molecular pathways involved in responses to thermal stress, we sampled families from a benign (20°C) and a stressful (24°C) thermal environment across the breeding cycle. While our previous work showed that *N. orbicollis* do not modify their behavior in response to temperature (Moss and Moore 2021), constitutive changes of gene expression of adults and larvae exposed to high temperatures across stages of care suggest that any phenotypic stabilization over this thermal range is at least partly due to the buffering action of background physiological processes. In parents, the predominant gene expression response to heat stress was downregulation, particularly of pathways involved in cuticle formation and carbohydrate metabolism (Supplementary Tables 12–13). Similarly, larvae showed downregulation of many pathways involved in cellular housekeeping functions (Supplementary Table 14). These findings are consistent with several long-term acclimation experiments reporting genome-wide downregulation in response to extreme heat stress, which could either point to an inability to mount systemic responses (i.e., due to insufficent evolutionary history of such extremes; Levine, Eckert, & Begun 2011; Becker et al., 2018) or an adaptive molecular mechanism for metabolic compensation (Yampolsky et al. 2014). The high temperature applied here was at the upper limit but still within the range experienced in the habitat our beetles were collected (Moss and Moore, 2021), suggesting the latter as a more likely explanation. In sum, thermal stress appears to be mitigated through the regulation of different functional pathways in parents vs larvae but follows the same general pattern – tempering all gene expression responses at high temperatures – which could impose carry-over effects on gene expression underlying parent-offspring and offspring-parent interactions

With our study, we provide a glimpse into these mechanisms by comparing the gene expression profiles of family members before and 24 hours after offspring colonize the brood ball. Thousands of differentially expressed genes were detected in larvae spanning many functional categories, some due to differences of sampling ages dictated by experimental design. While we cannot disentangle the direct effects being parented *per se* on gene expression (Capodeanu-Nägler et al., 2018; Mashoodh et al., 2021) from indirect effects associated with the rapid development and growth occurring at this early stage (Won et al., 2018), our sampling design was biologically realistic given that orphaned *N. orbicollis* larvae usually succumb to starvation within a day (Capodeanu-Nägler et al. 2018). In comparison, changes of parental gene expression associated with having larvae to parent were subtle (∼100–200 differentially expressed genes), in line with the hypothesis that the modification of behavior within a state involves a much smaller subset of the genes required to transition between states (i.e., *N. vespilloides* transitioning into parenting from a non-reproductive state differentially express up to 650 genes; Parker et al. 2015, Cunningham et al. 2017, Bell, Bukhari, & Sanogo, 2016; Benowitz, McKinney, Cunningham, et al., 2017; Cunningham et al., 2019).

As in previous work on the related *N. vespilloides* (Parker et al. 2015), males and females parenting together showed strong concordance in overall gene expression patterns even though, as has been found for every study of parenting in male burying beetles, we detected fewer significantly differentially expressed genes for males. More so than males, however, actively parenting females showed intriguing patterns of overlap (Fig 3B) and concordance (Fig 3G) with parented larvae in terms of differentially expressed genes and pathways (i.e., males showed anti-concordance with larvae; Fig 3H). Given that mothers in biparental pairs appear to perform a disproportionate share of larval provisioning (Moss & Moore, 2021), we propose that these overlapping pathways, which include lipid and carboxylic acid metabolism, could play a role in self-feeding (i.e., from a shared resource) and/or nutrient exchange (i.e., mouth-to-mouth regurgitation). In either case, genes with shared or reciprocal functional roles between family members could serve as important targets of selection during adaptation to stressful environments, which may demand more efficient performance of caregivers, offspring, or both.

Finally, we characterized patterns of gene expression in response to the combined stressors of parenting/being parented and thermal stress. If plasticity of gene expression is important for maintaining behavioral stability of families across environments, then genes underlying parent-offspring interactions should be expressed differently or include more/different genes dependent on environment. The first way we tested this was by comparing the magnitude of differential expression of parenting/being parented genes between 24°C and 20°C. We predicted that if plasticity of gene expression underlies the ability of parents to buffer environmental stress or for offspring to compensate independently, then responses would be more pronounced at the higher temperature. Instead, we found that genes for receiving care (i.e., in larvae) were differentially expressed to the same degree regardless of temperature and that genes for providing care (i.e., in males and females) showed less robust expression changes at 24°C than at 20°C (Fig 3A,B). Weaker differential expression at 24°C is nevertheless a form of plasticity, and like global gene expression responses to thermal stress, could reflect adaptive metabolic compensation by parents.

Another possibility is that plastic responses to parenting/being parented mediated by thermal stress involve completely different genes to those captured by the main effects of parenting. If true, then many differentially expressed genes should be captured by the interaction between parenting and temperature, and changes in expression of these genes before and during parenting should be more pronounced and/or change their sign at the higher temperature. Overall few genes satisfied this condition in any family member. While not entirely surprising given that interaction terms have lower statistical power than main effects, this suggests that individuals in the same behavioral state but at a different temperature still rely on more-or-less the same genetic program. The largest gene expression response occurred in heat stressed actively parenting mothers, and closer inspection of this ‘buffering’ gene set (n = 79) revealed the nature of this plasticity. Genes that were differentially expressed at 20°C (i.e., significantly increased in expression during parenting) did not change expression or a temporal shift followed by sustained expression (i.e., increased before parenting but then showed no further change during parenting; Fig 3C) at 24°C. Thus, while temperature may independently influence the expression of some parenting genes the effects are not synergistic, and rather than amplify differences between behavioral states, thermal stress appears to have a tempering effect.

Our inferences based on differential expression analysis were further corroborated by functional enrichment analysis of maternal buffering genes, which revealed significant downregulation of pathways involved in fatty acid oxidation and lipid processing (Supplementary Table 18). Lipids are the principal macronutrient used for fueling energetically intensive behavior in insects (e.g. flight: Canavoso et al. 2003), yet previous efforts to link post-hatching care (i.e., presumed to be the costliest form of care) to increased lipid metabolism in burying beetles found no relationship (Benowitz, McKinney, Roy-Zokan, Cunningham, & Moore, 2017). We suggest that females restrict the use of stored energy even more in an energetically demanding environment, implying that the combined stressors of temperature and parenting could directly mediate trade-offs between current and future reproduction. In line with this, actively parenting cichlids exposed to stressful levels of noise show pronounced gene expression changes in brain regions responsible for homeostatic functioning, suggesting that stressful environments may trigger a neural switch from offspring-promoting to self-promoting behaviors (Butler & Maruska 2021). Offspring of stressed mothers in our study in turn upregulated genes involved in forming the larval serum protein complex – a structure that sequesters amino acids early in development for the synthesis of adult proteins (Chrysanthis, Kaliafas, & Mintzas, 1994). Whether this gene expression response by larvae may help compensate for constrained parental investment or could simply be an indirect effect of plasticity in mothers remains to be seen. What is clear is that parental gene expression responses to larvae are tempered at higher temperatures but generally involve the same genes, meaning that the adaptive function most likely fulfilled by plasticity of gene expression is metabolic compensation. Thus, the scope for temperature adaptation to proceed *via* selection on alternative genetic programs underlying care would appear limited.

## ACKNOWLEDGEMENTS

We thank PJ Moore, EA Shelby, JT Washington, and KE Kollars for assistance with colony maintenance and Kathryn E. Kollars for donating her talents of illustration to create the figures in this manuscript. Robert Schmitz, Hannah Houston, James Parker, and Tyler Earp assisted with library preparation and Illumina sequencing. Eva Fischer provided helpful discussions and suggestions regarding statistical analyses and all members of the Fischer lab provided valuable and constructive feedback on this manuscript. We thank Romain Libbrecht, Tadeusz Kawecki and an anonymous reviewer for thorough and helpful peer reviews. The authors declare no conflicts of interest.

## DATA ACCESSIBILITY

All sequencing data is deposited in NCBI GenBank under the BioProject ID PRJA71530. Raw sequence reads are deposited in the Sequence Read Archive (SRA) under submission SUB9220525 and Accession numbers SAMN18349415– SAMN18349472 and the assembled transcriptome is deposited in the Transcriptome Shotgun Assembly (TSA) Sequence Database under submission SUB9312680. Supporting data and R scripts are deposited as an Open Science Framework project: osf.io/ztyqa.

## BENEFITS GENERATED

Benefits from this research accrue from the sharing of our data and results on public databases as described above.

## AUTHOR CONTRIBUTIONS

JBM and AJM conceived and designed the study. JBM, ECM, CBC, and AJM collected and analyzed the data. JBM, CBC, and AJM wrote the paper with input from ECM. All authors gave final approval for publication.

## Notes

https://www.osf.io/ztyqa

### Competing Interest Statement

The authors have declared no competing interest.

## References

Alexa, A., & Rahnenfuhrer, J. (2020). topGO: Enrichment analysis for gene ontology. R Package Version 2.42.0.

Alhendi, A. (2019). countToFPKM: Convert counts to fragments per kilobase of transcript per million (FPKM). R Package Version 3.1.0.

Becker, D., Reydelet, Y., Lopez, J. A., Jackson, C., Colbourne, J. K., Hawat, S., … Paul, R. J. (2018). The transcriptomic and proteomic responses of Daphnia pulex to changes in temperature and food supply comprise environment-specific and clone-specific elements. BMC Genomics, 19, 1–30. doi:10.1186/S12864-018-4742-6/FIGURES/15

Bell, A. M., Bukhari, S. A., & Sanogo, Y. O. (2016). Natural variation in brain gene expression profiles of aggressive and nonaggressive individual sticklebacks. Behaviour, 153, 1723–1743. doi:10.1163/1568539X-00003393

Benjamini, Y., & Hochberg, Y. (1995). Controlling the False Discovery Rate: a Practical and Powerful Approach to Multiple Testing. J. R. Statist. Soc. B (Vol. 57).

Benowitz, K. M., McKinney, E. C., Cunningham, C. B., & Moore, A. J. (2017). Relating quantitative variation within a behavior to variation in transcription. Evolution, 71, 1999–2009. doi:10.1111/evo.13273

Benowitz, K. M., McKinney, E. C., Roy-Zokan, E. M., Cunningham, C. B., & Moore, A. J. (2017). The role of lipid metabolism during parental care in two species of burying beetle (Nicrophorus spp.). Animal Behaviour, 129, 143–149. doi:10.1016/j.anbehav.2017.05.019

Benowitz, K. M., & Moore, A. J. (2016). Biparental care is predominant and beneficial to parents in the burying beetle Nicrophorus orbicollis (Coleoptera: Silphidae). Biological Journal of the Linnean Society, 119, 1082–1088. doi:10.1111/bij.12830

Bloch, N. I., Corral-López, A., Buechel, S. D., Kotrschal, A., Kolm, N., & Mank, J. E. (2018). Early neurogenomic response associated with variation in guppy female mate preference. Nature Ecology & Evolution 2018 2, 1772–1781. doi:10.1038/s41559-018-0682-4

Bolger, A. M., Lohse, M., & Usadel, B. (2014). Trimmomatic: A flexible trimmer for Illumina sequence data. Bioinformatics, 30, 2114–2120. doi:10.1093/bioinformatics/btu170

Boratyn, G. M., Thierry-Mieg, J., Thierry-Mieg, D., Busby, B., & Madden, T. L. (2019). Magic-BLAST, an accurate RNA-seq aligner for long and short reads. BMC Bioinformatics, 20, 405. doi:10.1186/s12859-019-2996-x

Bukhari, S. A., Saul, M. C., James, N., Bensky, M. K., Stein, L. R., Trapp, R., & Bell, A. M. (2019). Neurogenomic insights into paternal care and its relation to territorial aggression. Nature Communications, 10, 4437. doi:10.1038/s41467-019-12212-7

Cahill, K. M., Huo, Z., Tseng, G. C., Logan, R. W., & Seney, M. L. (2018). Improved identification of concordant and discordant gene expression signatures using an updated rank-rank hypergeometric overlap approach. Scientific Reports 8, 1–11. doi:10.1038/s41598-018-27903-2

Canavoso, L. E., Jouni, Z. E., Karnas, K. J., Pennington, J. E., & Wells, M. A. (2003). Fat metabolism in insects. Annual Review of Nutrition, 21, 23–46. doi:10.1146/ANNUREV.NUTR.21.1.23

Capodeanu-Nägler, A., Prang, M. A., Trumbo, S. T., Vogel, H., Eggert, A. K., Sakaluk, S. K., & Steiger, S. (2018). Offspring dependence on parental care and the role of parental transfer of oral fluids in burying beetles. Frontiers in Zoology, 15, 33. doi:10.1186/s12983-018-0278-5

Cheviron, Z. A., & Brumfield, R. T. (2011). Genomic insights into adaptation to high-altitude environments. Heredity 108, 354–361. doi:10.1038/hdy.2011.85

Cheviron, Z. A., Whitehead, A., & Brumfield, R. T. (2008). Transcriptomic variation and plasticity in rufous-collared sparrows (Zonotrichia capensis) along an altitudinal gradient. Molecular Ecology, 17, 4556–4569. doi:10.1111/J.1365-294X.2008.03942.X

Chrysanthis, G., Kaliafas, A. D., & Mintzas, A. C. (1994). Biosynthesis and tissue distribution of four major larval serum proteins during development of Ceratitis capitata (Diptera). Insect Biochemistry and Molecular Biology, 24, 811–818. doi:10.1016/0965-1748(94)90109-0

Clauser, A. J., & McRae, S. B. (2016). Plasticity in incubation behavior and shading by king rails Rallus elegans in response to temperature. Journal of Avian Biology, 48, 479–488. doi: 10.1111/jav.01056

Clutton-Brock, T. H. (1991). The evolution of parental care. Princeton, New Jersey, U.S.A.: Princeton University Press.

Cunningham, C. B., Badgett, M. J., Meagher, R. B., Orlando, R., & Moore, A. J. (2017). Ethological principles predict the neuropeptides co-opted to influence parenting. Nature Communications, 81, 4225. doi:10.1038/ncomms14225

Cunningham, C. B., Ji, L., McKinney, E. C., Benowitz, K. M., Schmitz, R. J., & Moore, A. J. (2019). Changes of gene expression but not cytosine methylation are associated with male parental care reflecting behavioural state, social context and individual flexibility. Journal of Experimental Biology, 222, 1–9. doi:10.1242/jeb.188649

Cunningham, C. B., Ji, L., Wiberg, R. A. W., Shelton, J., McKinney, E. C., Parker, D. J., … Moore, A. J. (2015). The genome and methylome of a beetle with complex social behavior, Nicrophorus vespilloides (coleoptera: Silphidae). Genome Biology and Evolution, 7, 3383–3396. doi:10.1093/gbe/evv194

Eberwine, J., & Kim, J. (2015). Cellular deconstruction: Finding meaning in individual cell variation. Trends in Cell Biology, 25, 569–578. doi:10.1016/J.TCB.2015.07.004

Eggert, A.-K., & Müller, J. K. (1997). Biparental care and social evolution in burying beetles: lessons from the larder. In J. C. Choe & B. I. Crespi (Eds.), The Evolution of Social Behavior in Insects and Arachnids (pp. 216–236). Cambridge, Massachusetts, USA: Cambridge University Press. doi:10.1017/cbo9780511721953.011

El-Gebali, S., Mistry, J., Bateman, A., Eddy, S. R., Luciani, A., Potter, S. C., … Finn, R. D. (2019). The Pfam protein families database in 2019. Nucleic Acids Research, 47, D427–D432. doi:10.1093/nar/gky995

Fischer, E. K., Roland, A. B., Moskowitz, N. A., Tapia, E. E., Summers, K., Coloma, L. A., & O’Connell, L. A. (2019). The neural basis of tadpole transport in poison frogs. Proceedings of the Royal Society B: Biological Sciences, 286, 20191084. doi:10.1098/rspb.2019.1084

Fischer, E. K., Hauber, M. E., & Bell, A. M. (2021). Back to the basics? Transcriptomics offers integrative insights into the role of space, time and the environment for gene expression and behaviour. Biology Letters, 17, 20210293. doi:10.1098/RSBL.2021.0293

Huerta-Cepas, J., Szklarczyk, D., Heller, D., Hernández-Plaza, A., Forslund, S. K., Cook, H., … Bork, P. (2019). EggNOG 5.0: A hierarchical, functionally and phylogenetically annotated orthology resource based on 5090 organisms and 2502 viruses. Nucleic Acids Research, 47, D309–D314. doi:10.1093/nar/gky1085

Jacobs, C. G. C., Steiger, S., Heckel, D. G., Wielsch, N., Vilcinskas, A., & Vogel, H. (2016). Sex, offspring and carcass determine antimicrobial peptide expression in the burying beetle. Scientific Reports 6, 1–8. doi:10.1038/srep25409

Kim, D., Langmead, B., & Salzberg, S. L. (2015). HISAT: A fast spliced aligner with low memory requirements. Nature Methods, 12, 357–360. doi:10.1038/nmeth.3317

Körner, M., Vogelweith, F., Libbrecht, R., Foitzik, S., Feldmeyer, B., & Meunier, J. (2020). Offspring reverse transcriptome responses to maternal deprivation when reared with pathogens in an insect with facultative family life. Proceedings of the Royal Society B, 287, 20200440. doi:10.1098/RSPB.2020.0440

Kriventseva, E. V., Kuznetsov, D., Tegenfeldt, F., Manni, M., Dias, R., Simão, F. A., & Zdobnov, E. M. (2019). OrthoDB v10: Sampling the diversity of animal, plant, fungal, protist, bacterial and viral genomes for evolutionary and functional annotations of orthologs. Nucleic Acids Research, 47, D807–D811. doi:10.1093/nar/gky1053

Levine, M. T., Eckert, M. L., & Begun, D. J. (2011). Whole-genome expression plasticity across tropical and temperatee Drosophila melanogaster populations from Eastern Australia. Molecular Biology and Evolution, 28, 249–256. doi.org/10.1093/molbev/msq197

Li, W., & Godzik, A. (2006). Cd-hit: a fast program for clustering and comparing large sets of protein or nucleotide sequences. Bioinformatics, 22, 1658–1659. doi:10.1093/bioinformatics/btl158

Love, M. I., Huber, W., & Anders, S. (2014). Moderated estimation of fold change and dispersion for RNA-seq data with DESeq2. Genome Biology, 15, 550. doi:10.1186/s13059-014-0550-8

Butler, J. M., & Maruska, K. P. (2021). Noise during mouthbrooding impairs maternal care behaviors and juvenile development and alters brain transcriptomes in the African cichlid fish Astatotilapia burtoni. Genes, Brain and Behavior, 20, e12692. doi: 10.1111/gbb.12692

Mashoodh, R., Westoby, J., & Kilner, R. (2021). Evolved changes in DNA methylation in response to the sustained loss of parental care in the burying beetle. BioRxiv. doi:10.1101/2021.03.25.436923

McClintock, M. E., Hepp, G. R., & Kennamer, R. A. (2014). Plasticity of incubation behaviors helps Wood Ducks (Aix sponsa) maintain an optimal thermal environment for developing embryos. The Auk, 131, 672–680. doi:10.1642/AUK-14-57.1

Moss, J. B., & Moore, A. J. (2021). Constrained flexibility of parental cooperation limits adaptive responses to harsh conditions. Evolution, 75, 1835–1849. doi:10.1111/EVO.14285

Nijhout, H. F., Best, J. A., & Reed, M. C. (2019). Systems biology of robustness and homeostatic mechanisms. Wiley Interdisciplinary Reviews: Systems Biology and Medicine, 11, e1440. doi:10.1002/WSBM.1440

Oldekop, J. A., Smiseth, P. T., Piggins, H. D., & Moore, A. J. (2007). Adaptive switch from infanticide to parental care: How do beetles time their behaviour? Journal of Evolutionary Biology, 20, 1998–2004. doi:10.1111/J.1420-9101.2007.01364.X

Oleksiak, M. F., Roach, J. L., & Crawford, D. L. (2005). Natural variation in cardiac metabolism and gene expression in Fundulus heteroclitus. Nature Genetics, 37, 67–72. doi: 10.1038/ng1483

Palmer, W. J., Duarte, A., Schrader, M., Day, J. P., Kilner, R., & Jiggins, F. M. (2016). A gene associated with social immunity in the burying beetle Nicrophorus vespilloides. Proceedings of the Royal Society B: Biological Sciences, 283(1823), 20152733. doi:10.1098/rspb.2015.2733

Parker, D. J., Cunningham, C. B., Walling, C. A., Stamper, C. E., Head, M. L., Roy-Zokan, E. M., … Moore, A. J. (2015). Transcriptomes of parents identify parenting strategies and sexual conflict in a subsocial beetle. Nature Communications, 6, 8449. doi:10.1038/ncomms9449

Peck, L. S., Thorne, M. A. S., Hoffman, J. I., Morley, S. A., & Clark, M. S. (2015). Variability among individuals is generated at the gene expression level. Ecology, 96, 2004–2014. doi: 10.1890/14-0726.1.

Pertea, M., Pertea, G. M., Antonescu, C. M., Chang, T. C., Mendell, J. T., & Salzberg, S. L. (2015). StringTie enables improved reconstruction of a transcriptome from RNA-seq reads. Nature Biotechnology, 33, 290–295. doi:10.1038/nbt.3122

R Core Development Team. (2019). R: A language and environment for statistical computing. R Foundation for Statistical Computing.

Ray, S., Tzeng, R. Y., DiCarlo, L. M., Bundy, J. L., Vied, C., Tyson, G., Nowakowski, R., & Arbeitman, M. N. (2016). An examination of dynamic gene expression changes in the mouse brain during pregnancy and the postpartum period. G2, 6, 221–233. doi: 10.1534/g3.115.020982

Rivera, H. E., Aichelman, H. E., Fifer, J. E., Kriefall, N. G., Wuitchik, D. M., Wuitchik, S. J. S., & Davies, S. W. (2021). A framework for understanding gene expression plasticity and its influence on stress tolerance. Molecular Ecology, 30, 1381–1397. doi:10.1111/MEC.15820

Royle, N. J., Smiseth, P. T., & Kölliker, M. (2012). The Evolution of Parental Care. Oxford, U.K.: Oxford University Press.

Scott, M. P. (1998). The ecology and behavior of burying beetles. Annual Review of Entomology, 43, 595–618. doi:10.1146/annurev.ento.43.1.595

Scott, M. P., & Traniello, J. F. A. (1990). Behavioural and ecological correlates of male and female parental care and reproductive success in burying beetles (Nicrophorus spp.). Animal Behaviour, 39, 274–283. doi:10.1016/S0003-3472(05)80871-1

Sharpe, L. L., Bayter, C., & Gardner, J. L. (2021). Too hot to handle? Behavioural plasticity during incubation in a small, Australian passerine. Journal of Thermal Biology, 98, 102921. doi:10.1016/J.JTHERBIO.2021.102921

Simão, F. A., Waterhouse, R. M., Ioannidis, P., Kriventseva, E. V., & Zdobnov, E. M. (2015). BUSCO: Assessing genome assembly and annotation completeness with single-copy orthologs. Bioinformatics, 31, 3210–3212. doi:10.1093/bioinformatics/btv351

Song, L., & Florea, L. (2015). Rcorrector: Efficient and accurate error correction for Illumina RNA-seq reads. GigaScience, 4(1), 48. doi:10.1186/s13742-015-0089-y

Trumbo, S. T. (1991). Reproductive benefits and the duration of paternal care in a biparental burying beetle, Nicrophorus orbicollis. Behaviour, 117, 82–105. doi:10.1163/156853991X00139

Vitting-Seerup, K., & Sandelin, A. (2019). IsoformSwitchAnalyzeR: analysis of changes in genome-wide patterns of alternative splicing and its functional consequences. Bioinformatics, 35(21), 4469–4471. doi:10.1093/bioinformatics/btz247

Walter, W., Sánchez-Cabo, F., & Ricote, M. (2015). GOplot: an R package for visually combining expression data with functional analysis. Bioinformatics, 31, 2912–2914. doi:10.1093/BIOINFORMATICS/BTV300

Wilson, E. O. (1975). Sociobiology: The New Synthesis. Cambridge, U.K.: Belknap.

Won, H. I., Schulze, T. T., Clement, E. J., Watson, G. F., Watson, S. M., Warner, R. C., … Davis, P. H. (2018). De novo assembly of the burying beetle Nicrophorus orbicollis (Coleoptera: Silphidae) transcriptome across developmental stages with identification of key immune transcripts. Journal of Genomics, 6, 41. doi:10.7150/JGEN.24228

Yampolsky, L. Y., Zeng, E., Lopez, J., Williams, P. J., Dick, K. B., Colbourne, J. K., & Pfrender, M. E. (2014). Functional genomics of acclimation and adaptation in response to thermal stress in Daphnia. BMC Genomics, 15, 859. doi: 10.1186/1471-2164-15-859

Ziadie, M. A., Ebot-Ojong, F., McKinney, E. C., & Moore, A. J. (2019). Evolution of personal and social immunity in the context of parental care. American Naturalist, 193, 296–308. doi:10.1086/701122

