## Supplementary Methods 1 for "Limited plasticity in gene expression for providing or receiving parental care under different temperatures"

Reads associated with NCBI BioProject PRJNA371654, which includes one 350 bp library and one 6.5k mate-pair library squeneced with 150 bp paired end reads, were reassembled. Programs were run with default parameters unless specified otherwise. Reads were QC’ed before and after cutadapt with FASTQC (v0.11.9; Andrews, 2010). cutadapt (v2.8; Martin, 2011) was run with parameters to trim read and remove adapter contamination: -a AGATCGGAAGAGC -A AGATCGGAAGAGC --pair-filter=any -b CTGTCTCTTATACACATCT -b AGATGTGTATAAGAGACAG -a GATCGGAAGAGCACACGTCTGAACTCCAGTCAC --trim-n -O 3 -u -2 -U -2 -q 15,15 -m 30. FLASh (v1.2.11; Magoc and Salzberg, 2011) were used to merge overlapped paired end reads from the 350 bp library. Platanus (v2.2.2; Kajitani et al., 2019) was used to assemble contigs (parameters: -k 25 -s 3 -m 500) with all files for the 350 bp library, to scaffold with the non-overlapped 350 bp mate-pair library and the 6.5k mate-pair library, and phase with the same files from scaffolding. We used Conterminator (vGenome Biology Release; Steinegger and Salzberg, 2020) to remove contamination from identified sources from our previous work; *Homo sapiens*, *Morganella morganii*, *Caenorhabditis elegans*, *Yarrowia lipolytica*, and *Escherichia coli*. We used a BUSCO analysis (v4.0.5; --augustus_species tribolium2012; Simão et al., 2015) using the endopterygota_odb10 dataset to assess gene compliment completeness and BBMap’s stats.sh script to summarize the assembly metrics (v38.90; Bushnell, 2014). All analyses were conducted on the University of Georgia’s Center for Research Computing Sapelo2 computing cluster.

Results

The reassembly resulted in a more complete, but more fragmented assembly. The genome assembly totaled 242.4 Mb in 31,894 scaffolds, had a N50 of 1884, had an L50 of 27.3 kb, and a max scaffold length of 740.1 kb. The BUSCO analysis returned results of Complete: 96.1% [Single Copy: 94.3%, Duplicated: 1.8%], Fragmented :2.1%, Missing: 1.8% orthologs for the 2,1224 HMM’s searched.

Reference:

Andrews, S. (2010). FastQC: A Quality Control Tool for High Throughput Sequence Data. Available online at: http://www.bioinformatics.babraham.ac.uk/projects/fastqc/

Bushnell B. 2014. BBMap: A Fast, Accurate, Splice-Aware Aligner. Available online at: https://www.osti.gov/biblio/1241166-bbmap-fast-accurate-splice-aware-aligner.

Kajitani R, Yoshimura D, Okuno M, Minakuchi Y, Kagoshima H, Fujiyama A, Kubokawa K, Kohara Y, Toyoda A, Itoh T. 2019. Platanus-allee is a de novo haplotype assembler enabling a comprehensive access to divergent heterozygous regions. Nature Communications 10, 1702.

Magoc T, Salzberg S. 2011. FLASH: Fast length adjustment of short reads to improve genome assemblies. Bioinformatics 27, 2957-63.

Martin M. 2011. Cutadapt removes adapter sequences from high-throughput sequencing reads.

EMBnet.journal, 17,10-12.

Simão FA, Waterhouse RM, Ioannidis P, Kriventseva EV, Zdobnov EM. 2015. BUSCO: assessing genome assembly and annotation completeness with single-copy orthologs. Bioinformatics 31, 3210-3212.

Steinegger M, Salzberg SL. 2020. Terminating contamination: large-scale search identifies more than 2,000,000 contaminated entries in GenBank. Genome Biology 21, 115.
